# Na_v_1.5 regulates intrinsic cardiac neuronal excitability and expands the cellular basis of *SCN5A* channelopathies

**DOI:** 10.64898/2026.10.01.755938

**Authors:** A. Jean, H. Rabeony, C. Magaud, M. Derangeon, A. Tessier, B. Gellen, J.-F. Faivre, F. Charpentier, A. Chatelier

## Abstract

**Background:** Loss- and gain-of-function variants in *SCN5A*, encoding the cardiac sodium channel Na_v_1.5, are major causes of inherited cardiac channelopathies. Their pathophysiology has almost exclusively been attributed to altered cardiomyocyte excitability and cardiac conduction. However, growing evidence indicates that autonomic dysfunction contributes to these disorders, raising the possibility that Na_v_1.5 dysfunction may also affect the intrinsic cardiac nervous system.

**Objectives:** We investigated whether Na_v_1.5 is functionally expressed in murine intrinsic cardiac neurons and evaluated its contribution to neuronal excitability.

**Methods:** *Scn5a* expression was investigated by digital PCR and immunocytochemistry in isolated murine intracardiac neurons. Whole-cell patch-clamp recordings were used to characterize sodium currents and action potential properties in wild-type and *Scn5a*^*+/−*^ mice.

**Results:** *Scn5a* transcripts and Na_v_1.5 protein were detected in intrinsic cardiac neurons. Voltage-clamp recordings identified a tetrodotoxin-resistant sodium current accounting for approximately 8% of the total sodium current and displaying pharmacological and biophysical properties consistent with Na_v_1.5. Activation and inactivation analyses indicated that Na_v_1.5 is the predominant sodium conductance available at most polarized membrane potentials. Accordingly, *Scn5a* haploinsufficiency selectively altered action potential threshold, maximal upstroke velocity, overshoot and duration in neurons resting below −60 mV, whereas more depolarized neurons were essentially unaffected. No structural or neurochemical remodeling of intrinsic cardiac ganglia was detected.

**Conclusion:** Na_v_1.5 is a functional determinant of intrinsic cardiac neuronal excitability. These findings identify the intrinsic cardiac nervous system as a new cellular target of *SCN5A* dysfunction and suggest that inherited cardiac sodium channelopathies could involve alterations in both myocardial and intrinsic neuronal electrophysiology.

## Introduction

Inherited cardiac sodium channelopathies caused by variants in *SCN5A* are among the most extensively studied inherited arrhythmia syndromes. Loss- and gain-of-function mutations of *SCN5A*, encoding the voltage-gated sodium channel (VGSC) Na_v_1.5, underlie a broad spectrum of disorders including Brugada syndrome, long QT syndrome type 3, progressive cardiac conduction disease and familial atrial fibrillation^1,2^. Because Na_v_1.5 is the predominant sodium channel in cardiomyocytes, the pathophysiology of these diseases has largely been interpreted through alterations in myocardial excitability, impulse conduction and cardiac electrical remodeling. However, despite major advances in our understanding of Na_v_1.5 biology, cardiomyocyte dysfunction alone does not fully explain the marked variability in disease penetrance, arrhythmia susceptibility and clinical severity observed among mutation carriers.

Autonomic regulation contributes importantly to both inherited and acquired cardiac arrhythmias. Alterations in sympathovagal balance have been reported in Brugada syndrome, long QT syndrome and atrial fibrillation, and modulation of cardiac autonomic activity has emerged as a promising therapeutic strategy in several arrhythmogenic disorders^3,4^. Within the autonomic nervous system, the intrinsic cardiac nervous system (ICNS) occupies a unique position. Rather than acting as a simple relay between extracardiac autonomic inputs and the myocardium, the ICNS is now recognized as an integrative neuronal network with various electrophysiological properties capable of locally processing sensory information and dynamically regulating cardiac chronotropy, dromotropy and inotropy^5–9^. Experimental and clinical studies increasingly implicate this neuronal network in the initiation and maintenance of both atrial and ventricular arrhythmias^10–12^.

The ionic mechanisms governing intrinsic cardiac neuronal excitability remain, however, incompletely understood^13^. In particular, the molecular identity of VGSC responsible for action potential generation in intrinsic cardiac neurons remains largely unresolved. Together with tetrodotoxin-sensitive (TTX-S) channels, which have been identified at physiological scales, previous electrophysiological studies reported the presence of a tetrodotoxin-resistant (TTX-R) sodium current in canine intracardiac neurons, suggesting that Na_v_1.5 is expressed in the ICNS^14^. Nevertheless, direct evidence supporting Na_v_1.5 physiological contribution to intrinsic cardiac neuronal excitability is still lacking. Addressing this question is of particular importance because demonstration of functional Na_v_1.5 expression in intrinsic cardiac neurons would considerably broaden the current view of *SCN5A*-related channelopathies. Rather than exclusively affecting cardiomyocytes, *SCN5A* mutations could simultaneously alter the electrical properties of two tightly interconnected excitable tissues: the myocardium and its intrinsic neuronal network.

Accordingly, the aim of the present study was to determine whether Na_v_1.5 is functionally expressed in murine intrinsic cardiac neurons and to establish its contribution to neuronal excitability. Using complementary molecular, pharmacological and electrophysiological approaches, we identify Na_v_1.5 as the major TTX-R sodium channel in intrinsic cardiac neurons and demonstrate that it selectively regulates the excitability of hyperpolarized neuronal populations. These findings reveal the intrinsic cardiac nervous system as a new cellular target of *SCN5A* dysfunction and provide a new framework for understanding inherited cardiac sodium channelopathies.

## Methods

For more details, see the Supplemental Material.

### Animals

The research reported in this paper adhered to Animal Research: Reporting of In Vivo Experiments and Guide for the Care and Use of Laboratory Animals guidelines. All animal procedures were performed using adult C57/BL6 mice (8-20 weeks) of either sex in accordance to the guidelines from Directive 2010/63/EU of the European Parliament on the protection of animals used for scientific purposes. *Scn5a*^+/−^ mice and WT littermates were raised at the animal facility of Nantes University Health Research Institute (UTE – IRS-UN) for isolation and investigation of intrinsic cardiac neurons.

### Neuron dissociation

Isolation of murine intrinsic cardiac neurons was performed as previously described^15,16^. Briefly, mice were anesthetized with ketamine/xylazine and heparinized. Hearts were excised, and atrial area containing ganglionated regions were dissected, enzymatically digested, mechanically dissociated, and plated on laminin-coated 35-mm dishes.

### Immunocytochemistry and immunohistochemistry

Freshly isolated neurons or heart cryosections were fixed, permeabilized, blocked, and incubated with primary and secondary antibodies (see Supplementary Methods). Isolated neurons were labeled for βIII-tubulin/TUJ1 and Na_v_1.5. Cryosections were labeled for choline acetyltransferase (ChAT) and tyrosine hydroxylase (TH). Nuclei were counterstained with DAPI. Images were acquired using an FV3000 confocal laser scanning microscope (Olympus). Quantification was performed by manual cell counting using ImageJ (NIH, Bethesda, MD, USA). Antibodies are listed in Supplementary Table S1.

### Electrophysiology

Whole-cell patch-clamp recordings were performed at room temperature within 30 hours of neuronal dissociation. Data were acquired using an Axopatch 200B amplifier, low-pass filtered at 5 kHz, digitized at 20 kHz with a Digidata 1550B interface, and analyzed using pCLAMP 11 (Molecular Devices, San Jose, CA, USA). For detailed solutions and protocols, see Supplementary Methods.

Voltage-gated sodium currents were recorded in voltage-clamp mode from a holding potential of −120 mV. Pharmacological sensitivity was assessed using tetrodotoxin (TTX; 300 nM and 2 μM), A-803467 (0.5 μM), and extracellular sodium replacement by N-methyl-D-glucamine (NMDG). Current-voltage relationships, activation curves, and steady-state inactivation curves were obtained using standard protocols. TTX-R currents were recorded in 300 nM TTX, and TTX-S currents were calculated by digital subtraction.

Action potentials (AP) were recorded in current-clamp mode during depolarizing current injections of increasing amplitude (20-pA increments, 500 ms). Rheobase was defined as the minimal current required to elicit an action potential. Action potential threshold, overshoot, maximal upstroke velocity (dV/dtmax), action potential duration at 50% repolarization (APD50), and afterhyperpolarization were measured using the Clampfit Advanced Analysis Module.

### RNA extraction, reverse transcription, and digital PCR

RNA was extracted from either microdissected atrial ganglionated regions or isolated neuronal samples. Isolated neuronal samples consisted of four independent pools of 10 neurons collected with patch pipettes. Reverse transcription and digital PCR were performed using a *QuantStudio™ Absolute Q™ Digital PCR System* (Thermo Fisher Scientific/Applied Biosystems, Supplementary Methods). TaqMan assays were used for Scn5a, ChAT, Tnnt2, and Gapdh. Results were expressed as copies per microliter and normalized to Gapdh.

### Statistical analysis

Statistical analysis was performed with GraphPad Prism (GraphPad Software, San Diego, CA). Data are presented as mean ± standard error of the mean. Depending on the data, statistical analyses were performed with Mann-Whitney test, t-test or Fisher exact test.

## Results

### Na_v_1.5 is expressed in murine intrinsic cardiac neurons

To determine whether Na_v_1.5 is expressed in the murine intrinsic cardiac nervous system, we first investigated Scn5a expression at both the transcript and protein levels. Digital PCR was performed on two complementary preparations: (i) the atrial ganglionated region, containing both cardiomyocytes and intrinsic cardiac neurons, and (ii) individually collected intracardiac neurons isolated by enzymatic dissociation (Figure 1A). The specificity of neuronal preparations was first verified using cell-specific markers. As expected, Tnnt2, a cardiomyocyte marker, was readily detected in the ganglionated tissue but was undetectable in isolated neuronal preparations, confirming the absence of cardiomyocyte contamination (Figure 1A). Conversely, expression of the cholinergic neuronal marker ChAT was enriched in isolated neurons, showing an approximately 450-fold increase relative to the ganglionated region after normalization to Gapdh. Importantly, *Scn5a* transcripts were detected in both preparations, demonstrating that intrinsic cardiac neurons express the gene encoding Na_v_1.5. To determine whether this transcript is translated into protein, Na_v_1.5 expression was further examined by immunocytochemistry. Na_v_1.5 immunoreactivity was observed in βIII-tubulin-positive neurons, with staining predominantly localized along the plasma membrane surrounding neuronal somata (Figure 1B).

**Figure 1.**
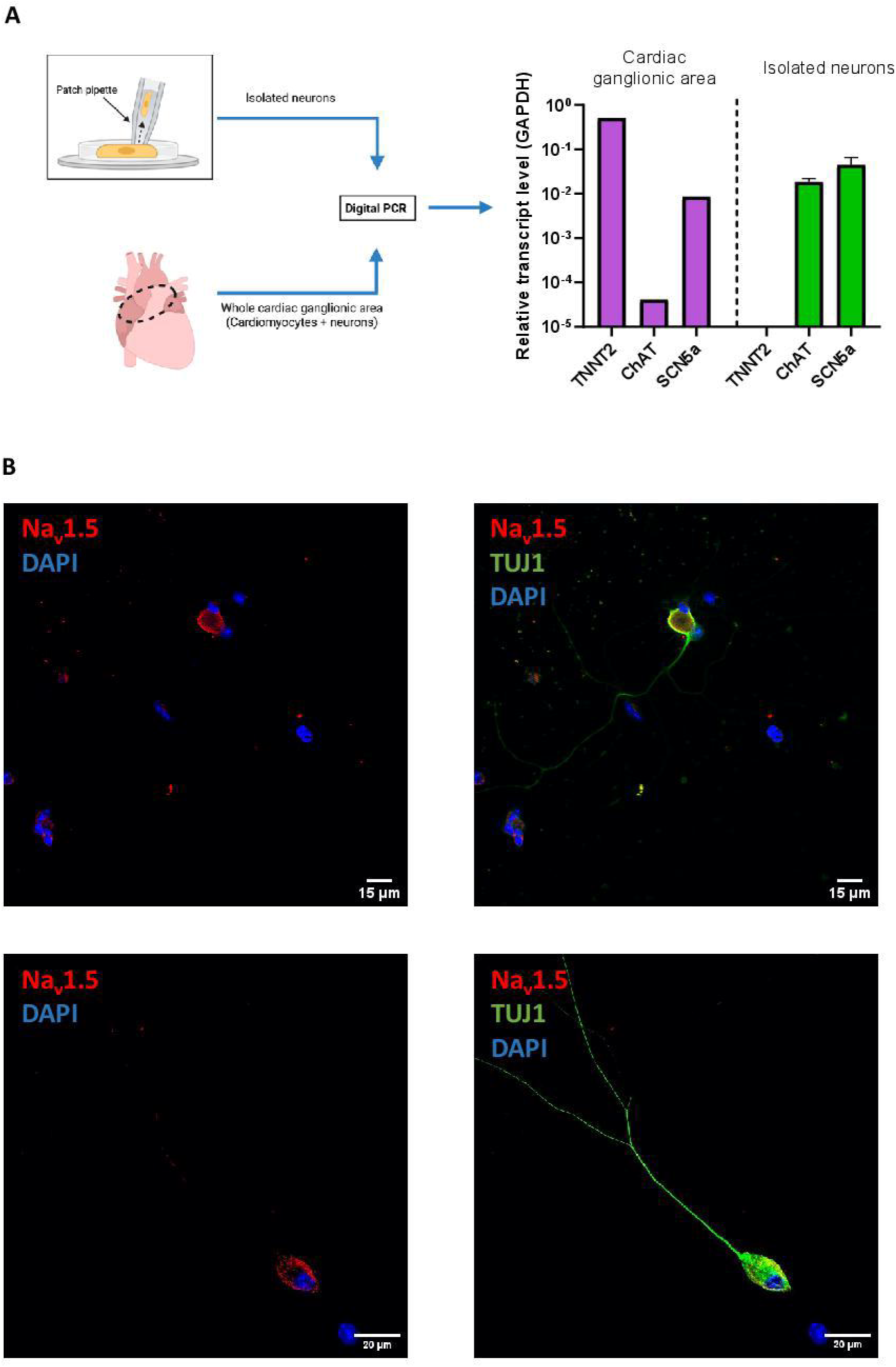
Expression of Scn5a transcripts and Na_v_1.5 in cardiac neurons. **A**. Left: Experimental workflow used for digital PCR analysis. Samples were obtained either from the whole atrial ganglionated region, containing intrinsic cardiac neurons and surrounding cardiac tissue, or from isolated intracardiac neurons harvested after enzymatic dissociation. Isolated neuronal samples consisted of four independent pools of 10 neurons collected individually with a patch-clamp pipette, corresponding to a total of 40 neurons analyzed. Right: Digital PCR was used to quantify the expression of Tnnt2, ChAT and *Scn5a*. Tnnt2 was used as a cardiomyocyte marker, ChAT as a cholinergic neuronal marker and Scn5a as the gene encoding Na_v_1.5. Primers were validated by digital PCR performed on the whole cardiac ganglionic area containing neurons and cardiomyocytes. **B**. Representative immunocytochemistry images of isolated intrinsic cardiac neurons labelled for βIII-tubulin/TUJ1 (green), Na_v_1.5 (red) and DAPI (blue). Na_v_1.5 immunoreactivity was detected around βIII-tubulin-positive neuronal somata.

**Together, these findings demonstrate that Na**_**v**_**1.5 is expressed at both the mRNA and protein levels in murine intrinsic cardiac neurons, supporting the hypothesis that this sodium channel may contribute directly to cardiac neuronal electrophysiology**.

### Na_v_1.5 constitutes the major tetrodotoxin-resistant sodium current in intrinsic cardiac neurons and is preferentially available at hyperpolarized membrane potentials

We next examined whether this channel contributes functionally to sodium current generation. Depolarizing voltage steps from a holding potential of −120 mV to −10 mV elicited large, rapidly activating and inactivating inward currents characteristic of VGSC (Figure 2A). Application of 300 nM TTX, which selectively blocks TTX-S sodium channels, markedly reduced the peak sodium current while consistently revealing a residual TTX-R component representing 8.4 ± 1.1% of the total current (n = 12; Figure 2A, B).

**Figure 2.**
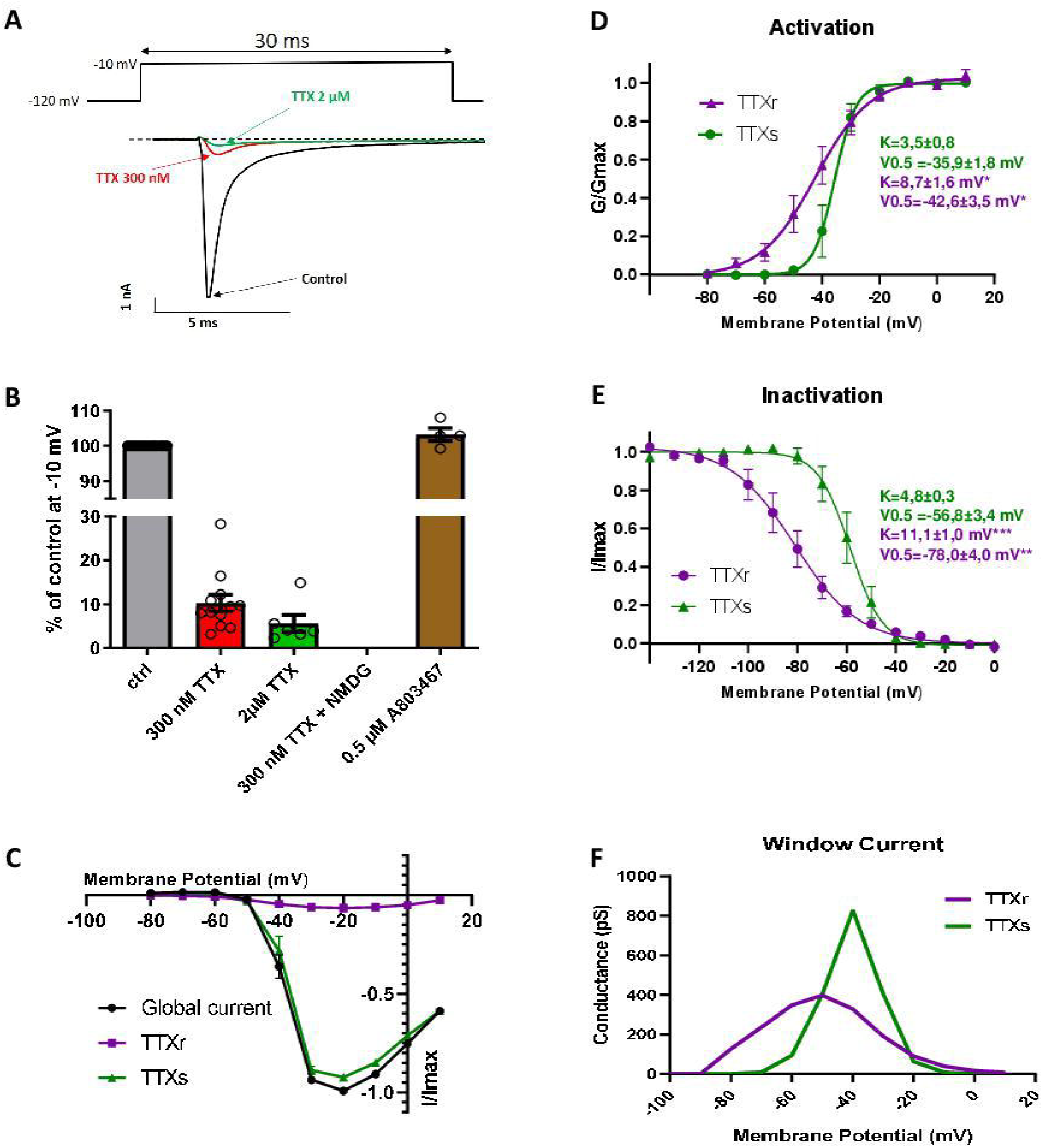
Na_v_1.5 constitutes the major tetrodotoxin-resistant sodium current in murine intrinsic cardiac neurons. **A**. Representative traces of the effect of TTX on peak sodium current (I_Na_) recorded at −10 mV. **B**. Impact of 300 nM (n=12) and 2 µM of TTX (n=6), of Na^+^ substitution by NMDG (n=4) and of 0.5 µM (n=4) of A803467, a specific blocker of Na_v_1.8, on peak current recorded at −10 mV. Data are expressed as percentage of the control current recorded in the same cell. **C**. Current–voltage relationships of total (Global) I_Na_ under control conditions (n = 6), TTX-resistant current recorded in the presence of 300 nM TTX (TTX-r; n = 5) and TTX-sensitive current (TTXs = Global - TTX-r, n=5). Currents were normalized to the maximal peak current recorded in each cell. **D**. Voltage dependence of activation of TTX-resistant and TTX-sensitive sodium currents obtained from the current–voltage relationships shown in panel C. **E**. Steady-state inactivation curves of TTX-resistant and TTX-sensitive sodium currents obtained using a standard two-step inactivation protocol (see methods section). **F**. Calculated conductances of TTX-resistant and TTX-sensitive sodium window current components derived from the activation and inactivation parameters shown in panels D and E. Statistical significance is indicated as *P < 0.05, **P < 0.01 and ***P < 0.001

Increasing the TTX concentration to 2 μM reduced the residual current by approximately 45%, consistent with the reported pharmacological sensitivity of Na_v_1.5^14,17,18^. In contrast, the selective Na_v_1.8 inhibitor A803467 (0.5 μM) produced no detectable reduction in total sodium current (Figure 2B), arguing against a major contribution of Na_v_1.8 under our experimental conditions. Finally, complete replacement of extracellular sodium with NMDG abolished the remaining inward current, confirming that it was carried exclusively by sodium ions (Figure 2B).

**Collectively, these data identify Na_v_1.5 as the predominant tetrodotoxin-resistant sodium channel underlying inward sodium current in murine intrinsic cardiac neurons.**

We then compared the voltage-dependent properties of TTX-S and TTX-R sodium currents.

Both current components reached maximal amplitudes near −20 mV (Figure 2C). However, marked differences were observed in their voltage dependence. Boltzmann analysis showed that TTX-R currents activated at significantly more hyperpolarized potentials than TTX-S currents (V_0.5_ = −42.6 ± 3.5 mV versus −35.9 ± 1.8 mV, respectively; p < 0.05), with distinct slope factors (Figure 2D). Even larger differences were observed for steady-state inactivation. TTX-R currents inactivated at substantially more negative membrane potentials than TTX-S currents (V_0.5_ = −78.0 ± 4.0 mV versus −56.8 ± 3.4 mV, respectively; p < 0.01, Figure 2E), indicating distinct voltage-dependent availability of the two channel populations.

To estimate the contribution of each current over the physiological voltage range, activation and inactivation curves were combined to calculate the steady-state window current as described previously^17^ (Figure 2F). This analysis demonstrated that below approximately −60 mV, the majority of available sodium current was carried by the TTX-R component. In contrast, as membrane potential became more depolarized, TTX-S channels progressively became the dominant sodium conductance.

**Thus, although Na**_**v**_**1.5 represents only a minor fraction of total peak sodium current, its voltage-dependent availability predicts a predominant functional contribution to neuronal excitability at resting membrane potentials (RMP) more negative than −60 mV**.

### Na_v_1.5 preferentially regulates the excitability of hyperpolarized intrinsic cardiac neurons

To determine the physiological contribution of Na_v_1.5 to intrinsic cardiac neuronal excitability, passive and active membrane properties of intracardiac neurons were compared between wild-type (WT) and *Scn5a* happlo-insufficient (*Scn5a* heterozygous knockout) mice. *Scn5a*^*+/−*^ mice are considered as a model of both Brugada Syndrome^19,20^ and congenital cardiac conduction disease^21,22^. The electrophysiological characteristics of all recorded neurons using whole-cell current-clamp recordings are summarized in Supplementary Table S2.

Depolarizing current injections elicited action potentials in both WT and *Scn5a*^*+/−*^ neurons (Figure 3A). As previously described in murine intrinsic cardiac neurons^7,8^, two principal firing behaviours were observed: accommodating and phasic neurons. Although *Scn5a* haploinsufficiency tended to increase the proportion of phasic neurons (86% versus 67% in WT) and consequently reduce the proportion of accommodating neurons, this difference did not reach statistical significance (Figure 3A). When all neurons were analyzed together, no significant differences were observed in resting membrane potential, rheobase, action potential threshold, overshoot, dV/dt_max_ or APD_50_ (Figure 3B and Supplementary Table S2).

**Figure 3.**
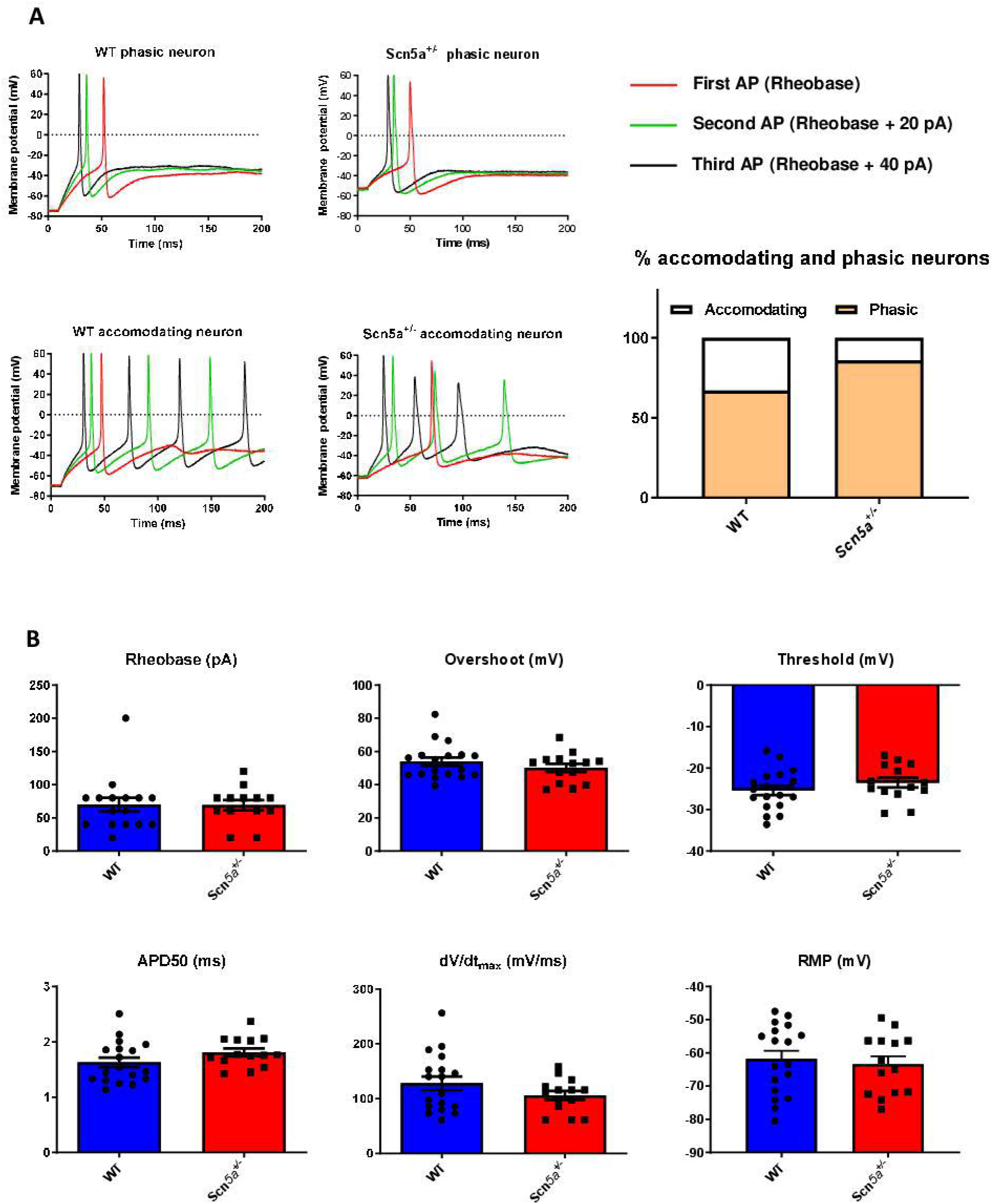
Global action potential properties are not significantly altered in *Scn5a*^*+/−*^ intrinsic cardiac neurons. **A**. Representative action potential recordings from WT and *Scn5a*^*+/*−^ intrinsic cardiac neurons displaying either accommodating or phasic firing patterns. Action potentials were elicited by depolarizing current injections of increasing amplitude (20-pA increments, 500-ms duration). The red trace corresponds to the first action potential elicited at rheobase, whereas the green and black traces correspond to subsequent current injections increased by 20 pA and 40 pA, respectively. The histogram shows the proportion of accommodating and phasic neurons in WT and *Scn5a*^*+/*−^ mice (WT, n = 18 neurons; *Scn5a*^*+/*−^, n = 14 neurons) **B**. Quantification of action potential properties in all recorded neurons, including rheobase, resting membrane potential (RMP), action potential overshoot, action potential threshold, action potential duration at 50% of repolarization (APD50), and maximal upstroke velocity (dV/dt_max_). No significant differences were detected between WT and *Scn5a*^*+/*−^ neurons when the entire neuronal population was analyzed.

Because window current analysis predicted that Na_v_1.5 availability depends strongly on resting membrane potential, neurons were stratified into two groups: RMP < −60mV and RMP > −60 mV (Figure 4A). Remarkably, this analysis uncovered a striking membrane potential-dependent phenotype. In neurons resting below −60 mV, *Scn5a* haploinsufficiency significantly reduced action potential overshoot (47.3 ± 2.9 *versus* 60.2 ± 3.7 mV in WT), decreased the maximal rate of depolarization (109.1 ± 13.2 *versus* 162.8 ± 16.1 mV/ms), depolarized the action potential threshold (−24.0 ± 1.8 versus −28.6 ± 1.1 mV) and prolonged APD50 (1.75 ± 0.08 *versus* 1.45 ± 0.08 ms) (Figure 4B, D). Rheobase was unchanged in this subgroup.

**Figure 4.**
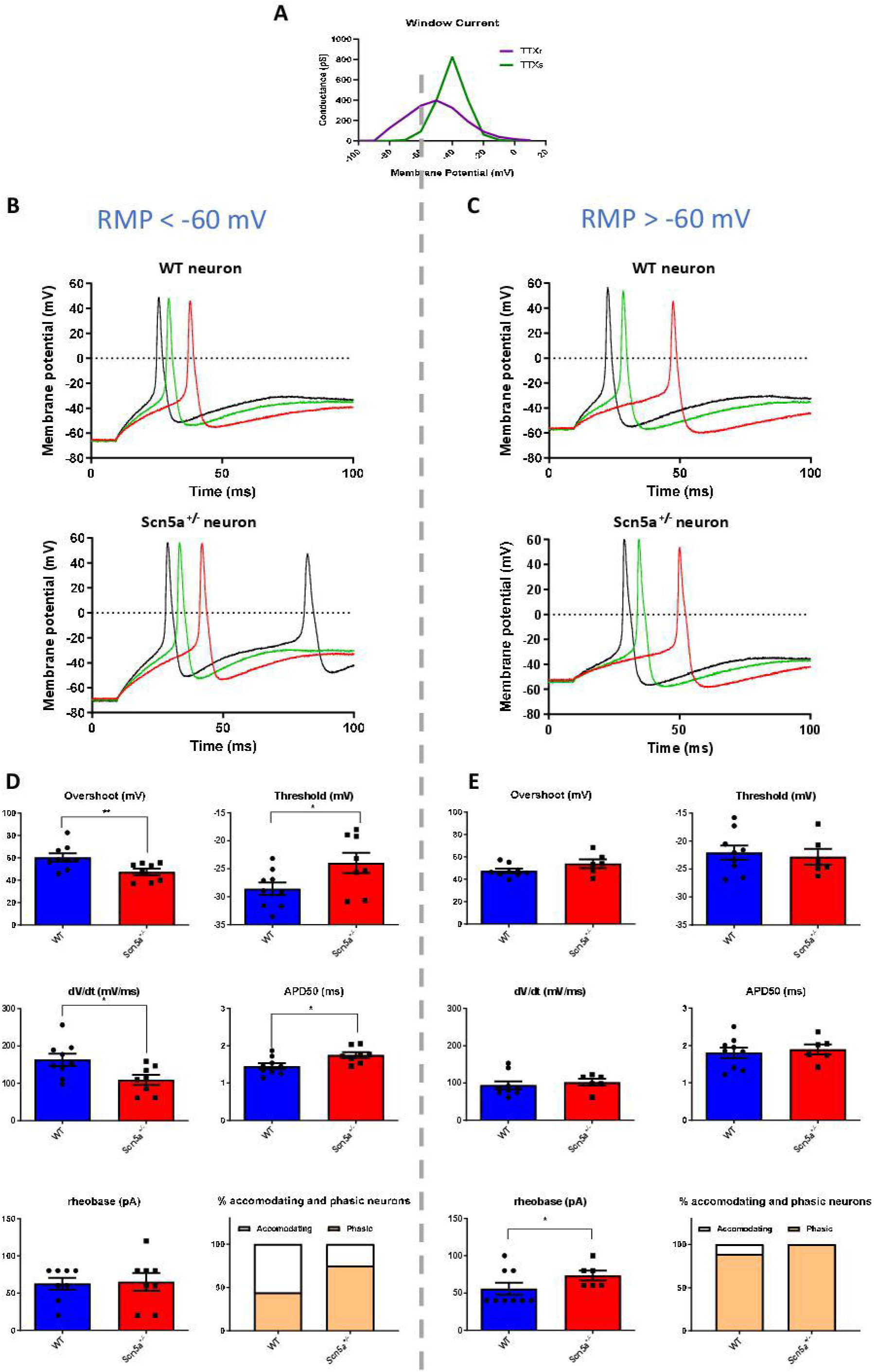
*Scn5a* haploinsufficiency selectively alters excitability of intrinsic cardiac neurons resting below −60 mV. **A**. Calculated TTX-resistant and TTX-sensitive window current derived from the activation and inactivation parameters shown in Figure 2. The −60 mV threshold was selected based on the predicted relative availability of TTX-resistant/Na_v_1.5 and TTX-sensitive sodium current components at resting membrane potentials. **B-C**. Representative action potential recordings from WT and *Scn5a*^*+/*−^ intrinsic cardiac neurons stratified according to resting membrane potential: RMP < −60 mV (B) and RMP > −60 mV (C). Action potentials were elicited by depolarizing current injections of increasing amplitude (20-pA increments, 500-ms duration). The red trace corresponds to the first action potential elicited at rheobase, whereas the green and black traces correspond to subsequent current injections increased by 20 pA and 40 pA, respectively. **D-E**. Quantification of action potential properties in neurons with RMP < −60 mV (D) and RMP > −60 mV (E). In neurons with RMP < −60 mV, *Scn5a* haploinsufficiency significantly decreased action potential overshoot and dV/dt_max_, depolarized the action potential threshold, and prolonged APD50. In contrast, neurons with RMP > −60 mV were largely unaffected, except for an increase in rheobase. RMP < −60 mV: WT, n = 9 neurons; *Scn5a*^*+/*−^, n = 8 neurons. RMP > −60 mV: WT, n = 9 neurons; *Scn5a*^*+/*−^, n = 6 neurons. Statistical significance is indicated as *P < 0.05 and **P < 0.01.

Conversely, neurons with RMP > −60 mV were largely insensitive to *Scn5a* haploinsufficiency. Neither action potential threshold, overshoot, dV/dt_max_, nor APD50 differed significantly between genotypes (Figure 4C, D). The only parameter affected was rheobase, which was significantly increased in *Scn5a*^*+/−*^ neurons compared with WT neurons (73.3 ± 6.7 versus 55.6 ± 8.0 pA) (Figure 4E). Interestingly, neuronal firing behavior also segregated according to resting membrane potential. Hyperpolarized neurons were predominantly phasic in *Scn5a*^*+/−*^ (75%) and accommodating in WT (56%)(56%) genotypes, whereas neurons resting above −60 mV were almost exclusively phasic (100% and 89%, respectively), consistent with our previous characterization of electrophysiological subtypes within the murine intrinsic cardiac nervous system.

**Overall, the selective alterations observed in hyperpolarized neurons closely match the voltage-dependent availability of Na**_**v**_**1.5, providing functional evidence that Na**_**v**_**1.5 preferentially regulates this population**.

### *Scn5a* haploinsufficiency does not induce structural remodeling of intrinsic cardiac ganglia

To determine whether the electrophysiological alterations observed in *Scn5a*^*+/−*^ neurons were associated with structural remodeling of the intrinsic cardiac nervous system, we next compared ganglion morphology and neuronal phenotype between WT and *Scn5a*^*+/−*^ mice labelled with choline acetyltransferase for cholinergic neurons and tyrosine hydroxylase for catecholaminergic neurons (Figure 5A). Quantitative morphometric analysis revealed no difference in neuronal soma area between genotypes (253.5 ± 7.3 μm^2^ in *Scn5a*^+/−^ versus 251.5 ± 12.1 μm^2^ in WT; Figure 5B). Likewise, the average number of neurons per ganglion was similar in both groups (120 ± 28.3 versus 90 ± 22.6 neurons per ganglion; Figure 5C). Finally, the proportion of tyrosine hydroxylase-positive neurons was unchanged (30.2 ± 4.7% in *Scn5a*^*+/−*^ versus 32.5 ± 6.2% in WT; Figure 5D).

**Figure 5.**
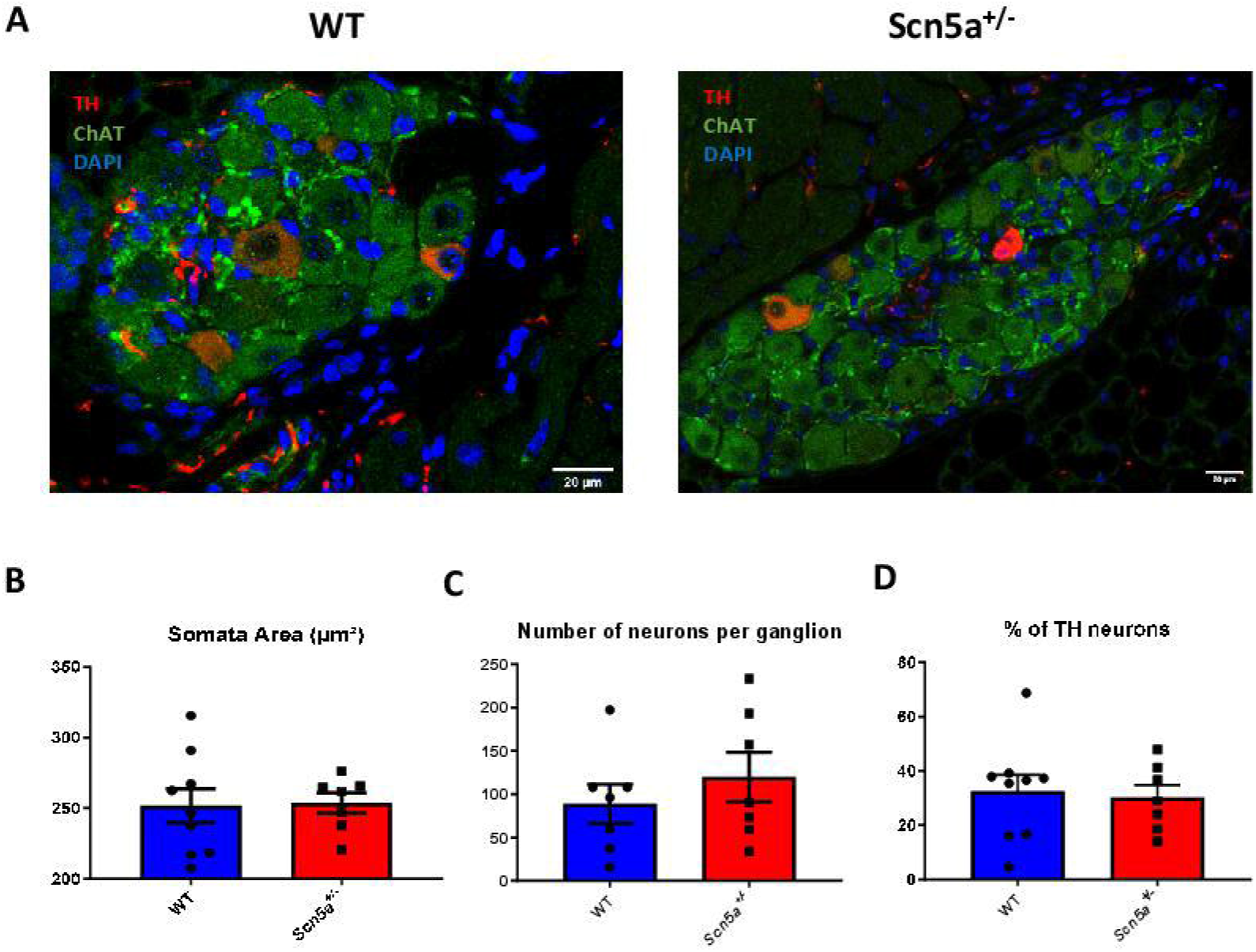
*Scn5a* haploinsufficiency does not induce detectable structural or neurochemical remodeling of intrinsic cardiac ganglia. **A**. Representative confocal images of intrinsic cardiac ganglia from WT and *Scn5a*^*+/*−^ mice immunolabelled for tyrosine hydroxylase (TH, red), choline acetyltransferase (ChAT, green), and DAPI (blue). **B**. Quantification of neuronal soma area in intrinsic cardiac ganglia from WT and *Scn5a*^*+/−*^ mice. **C**. Quantification of the average number of neurons per ganglion in *Scn5a*^*+/−*^ and WT mice. **D**. Quantification of the percentage of TH-positive neurons in WT and *Scn5a*^*+/−*^ intrinsic cardiac ganglia. No significant differences were detected between genotypes for neuronal soma area, neuronal number per ganglion, or proportion of TH-positive neurons. WT, n = 9 ganglia/mice; *Scn5a*^*+/*−^, n = 7 ganglia/mice, unless otherwise indicated.

**These observations indicate that *Scn5a* haploinsufficiency alters intrinsic cardiac neuronal excitability without inducing detectable structural or neurochemical remodeling of cardiac ganglia, supporting a primary electrophysiological mechanism**.

## Discussion

This study provides the first direct evidence that Na_v_1.5 is functionally expressed in mouse intrinsic cardiac neurons and contributes to the regulation of their excitability. By combining molecular, immunohistochemical, pharmacological and electrophysiological approaches, we demonstrate that (i) *Scn5a* transcripts and Na_v_1.5 protein are present in intrinsic cardiac neurons, (ii) Na_v_1.5 carries the major tetrodotoxin-resistant sodium current in these cells, and (iii) *Scn5a* haploinsufficiency alters action potential properties in a subset of neurons predicted to rely most heavily on Na_v_1.5. These findings broaden the cellular framework of *SCN5A*-related cardiac channelopathies beyond cardiomyocytes.

Voltage-gated sodium currents have been described previously in intrinsic cardiac neurons, but the molecular identity of the underlying channel isoforms remains poorly defined. Scornik et al. (2006)^14^ reported a TTX-resistant sodium current in canine intrinsic cardiac neurons and proposed Na_v_1.5 as its likely substrate. Our findings provide direct experimental support for this hypothesis. First, *Scn5a* transcripts were detected in isolated cardiac neurons devoid of cardiomyocyte contamination, while Na_v_1.5 protein was localized at the neuronal plasma membrane. Second, the residual sodium current remaining after application of 300 nM TTX was further inhibited by 2 µM TTX, a pharmacological profile consistent with the reported TTX sensitivity of Na_v_1.5 but not with the markedly lower TTX sensitivity of Na_v_1.8 or Na_v_1.9^14,17,18^. Finally, reducing *Scn5a* gene dosage modified action potential characteristics in the more negatively polarized neurons in a manner fully consistent with reduced sodium channel availability. Altogether, these independent lines of evidence establish that Na_v_1.5 is not simply expressed in intrinsic cardiac neurons but contributes functionally to their electrical activity.

Among tetrodotoxin-resistant isoforms, Na_v_1.8 has been proposed to contribute to intrinsic cardiac neuronal excitability^23^. However, the Na_v_1.8 selective inhibitor A803467 did not affect sodium currents in our experiments. This discrepancy may reflect differences in experimental conditions, because the selectivity of A803467 depends on membrane potential and concentration^24^. Although our data do not exclude Na_v_1.8 expression in specific neuronal subpopulations, they identify Na_v_1.5 as the principal TTX-resistant sodium channel in mouse intrinsic cardiac neurons under our recording conditions.

A key finding is that the consequences of *Scn5a* haploinsufficiency depend on neuronal RMP. Stratification by RMP revealed genotype-dependent alterations selectively in neurons resting below −60 mV, where Na_v_1.5 represents the predominant sodium conductance available for activation. Conversely, above −60 mV, genotype differences largely disappeared. This likely reflects reduced Na_v_1.5 availability in WT neurons at depolarized RMP, making their action potential properties closer to those of *Scn5a*^*+/−*^ neurons, in which Nav1.5 contribution is already reduced. Thus, the RMP-dependent phenotype strongly supports a functional role of Nav1.5 availability in shaping intrinsic cardiac neuronal excitability.

The identification of functional Na_v_1.5 expression in intrinsic cardiac neurons has important implications for understanding inherited cardiac sodium channelopathies. For more than two decades, the consequences of *SCN5A* mutations have largely been interpreted through their effects on cardiomyocyte excitability and impulse conduction. However, increasing evidence indicates that autonomic dysfunction contributes significantly to diseases such as Brugada syndrome, atrial fibrillation and long QT syndrome^3,4^, and the intrinsic cardiac nervous system has emerged as a critical integrative network controlling cardiac rhythm and arrhythmia susceptibility^10–12^. Our findings raise the possibility that *SCN5A* mutations simultaneously alter electrical activity in cardiomyocytes and intrinsic cardiac neurons. Such dual effects could contribute to the remarkable phenotypic variability observed among patients carrying *SCN5A* variants. In loss-of-function disorders such as Brugada syndrome, reduced Na_v_1.5 expression would be expected to decrease intrinsic cardiac neuronal excitability in addition to slowing myocardial conduction, whereas gain-of-function variants associated with atrial fibrillation or long QT syndrome type 3 may conversely enhance neuronal excitability. Whether these neuronal alterations directly contribute to arrhythmia initiation or maintenance remains to be established, but they introduce a previously underappreciated neuro-cardiac component to *SCN5A*-associated diseases. In this context, it will also be necessary to investigate the TTX-sensitive isoforms expressed within the ICNS and how these channels interact with Na_v_1.5 under physiological and pathological conditions.

Several limitations should nevertheless be acknowledged with regard to this study. First, our experiments were performed on isolated neurons, which do not fully reproduce the complex synaptic interactions occurring within intact intrinsic cardiac ganglia. Second, electrophysiological recordings were obtained under whole-cell patch-clamp conditions at room temperature. These experimental conditions do not preserve all physiological firing behaviors, particularly tonic discharge, which may be better investigated using intracellular microelectrode recordings at physiological temperature.

## Conclusion

In conclusion, this study identifies Na_v_1.5 as a functional determinant of intrinsic cardiac neuronal excitability and provides a new framework for understanding *SCN5A*-related cardiac channelopathies. Beyond its well-established role in cardiomyocytes, Na_v_1.5 should now be considered as a component of the intrinsic cardiac nervous system that may contribute to autonomic regulation of cardiac electrical activity. Considering the combined effects of *SCN5A* mutations on both cardiac muscle and cardiac neurons may therefore improve our understanding of inherited arrhythmias and open new therapeutic perspectives targeting the neuro-cardiac axis.

## List of abbreviations

AP: action potential
APD50: action potential duration at 50% repolarization
ChAT: choline acetyltransferase
ICNS: intrinsic cardiac nervous system
RMP: resting membrane potential
TH: tyrosine hydroxylase
TTX: tetrodotoxin
TTX-R: tetrodotoxin-resistant
TTX-S: tetrodotoxin-sensitive
VGSC: voltage-gated sodium channel
WT: wild-type

## Acknowledgements

This work has benefited from the facilities of ImageUP and PREBIOS platforms (University of Poitiers) and the technical assistance of Anne Cantereau and Isabelle Fixe. We also thank the staff from the animal facility of Nantes University Health Research Institute (UTE IRS-UN).

