## supplementary methods for "Na_v_1.5 regulates intrinsic cardiac neuronal excitability and expands the cellular basis of *SCN5A* channelopathies"

Jean A. *et al.*

#### Detailed isolation of murine intrinsic cardiac neurons

Mice (8-20 weeks) were intraperitoneally injected with heparin and anesthetized with xylazine (10 mg/kg) and ketamine (80 mg/kg). Following confirmation of the absence of nociceptive reflexes, hearts were rapidly excised and transferred into ice-cold Hanks' Balanced Salt Solution (HBSS). The fat pads area located between the two atria were dissected, cut into small pieces, and digested in 2 mL HBSS containing type II collagenase (3 mg/mL; Worthington) and dispase II (7.5 mg/mL; Gibco) for 30 min at 37°C. A second digestion was performed in 2 mL 0.25% trypsin-EDTA for 36 min at 37°C. After two washes in culture medium, cells were gently dissociated by trituration using fire-polished Pasteur pipettes pre-coated with fetal bovine serum (Eurobio Scientific) and plated onto laminin-coated 35-mm Petri dishes.

#### Immunocytochemistry

Freshly isolated neurons were fixed in 4% paraformaldehyde at 4°C. Samples were rinsed three times with phosphate-buffered saline (PBS), permeabilized for 10 min in PBS containing 0.1% Triton X-100 and 0.1% Tween-20, and blocked for 30 min at room temperature in PBS containing 1% bovine serum albumin (BSA). Neurons were incubated overnight at 4°C with primary antibodies and for 2 h at room temperature with secondary antibodies. Nuclei were stained with DAPI, and samples were mounted in Mowiol mounting medium.

#### Immunohistochemistry

Immunohistochemistry was performed on cryosections from paraformaldehyde-fixed heart samples. Hearts were cryoprotected overnight at 4°C in 30% sucrose, embedded in OCT compound (Tissue-Tek, Sakura Finetek, VWR), and frozen. Serial cryosections (50 μm) were obtained using a cryostat and mounted onto SuperFrost Plus adhesion slides. Sections were rinsed three times with PBS, incubated for 2 h at room temperature in blocking solution (PBS containing 1% BSA and 0.5% Triton X-100), incubated overnight at 4°C with primary antibodies, and then incubated for 3 h at room temperature with secondary antibodies. After DAPI staining, sections were mounted in Mowiol mounting medium.

**Voltage-clamp recordings**

Patch pipettes for sodium current recordings had a resistance of approximately 2 MΩ and were filled with the following intracellular solution (in mM): 140 CsF, 10 NaCl, 1 MgCl2, 1 EGTA, and 10 HEPES (pH 7.2 with CsOH). The extracellular solution contained (in mM): 150 NaCl, 5 KCl, 1 MgCl2, 10 glucose, and 10 HEPES (pH 7.4 with NaOH). For sodium-free experiments, NaCl was replaced by 150 mM N-methyl-D-glucamine (NMDG; pH adjusted with KOH). The liquid junction potential (+8.4 mV) was corrected before recordings. Linear leak and capacitive currents were removed using P/N leak subtraction.

Pharmacological sensitivity to TTX, NMDG, and A-803467 was assessed using depolarizing test pulses from a holding potential of -120 mV to -10 mV for 50 ms. Drugs and solutions were applied using a gravity-driven perfusion system (ALA Scientific Instruments). Current-voltage relationships were obtained from pre-pulse currents during the inactivation protocol. Peak inward currents were normalized to the maximal response of each cell.

The voltage dependence of activation was determined from relative membrane conductance using GNa = INa/(Vm - Vrev), where INa is the peak sodium current at test potential Vm and Vrev is the estimated reversal potential obtained by extrapolation of the current-voltage relationship.

Conductance values were normalized to maximal conductance and fitted with the Boltzmann equation $G/G_{max} = 1/\{1 + \exp[(V - V_{1/2})/k]\}$.

Steady-state inactivation was determined by measuring peak sodium current during a 20-ms test pulse to -10 mV following 500-ms conditioning pre-pulses from -120 mV to +10 mV from a holding potential of -120 mV. Peak currents were normalized to maximal current and fitted with a Boltzmann equation. TTX-resistant current-voltage, activation, and inactivation curves were obtained after perfusion of 300 nM TTX in the same cells. TTX-sensitive currents were calculated by subtracting TTX-resistant currents from total currents.

**Current-clamp recordings**

Patch pipettes for action potential recordings had a resistance of approximately 4 MΩ and were filled with the following intracellular solution (in mM): 130 K-gluconate, 10 KCl, 1 $MgCl_2$, 10 HEPES, 1 $CaCl_2$, 5 EGTA, 2 Mg-ATP, 10 $Na_2$-phosphocreatine, and 0.3 Na-GTP (pH 7.2 with KOH). The bath solution contained (in mM): 150 NaCl, 5 KCl, 1 $MgCl_2$, 1.5 $CaCl_2$, 10 glucose, and 10 HEPES (pH 7.4 with NaOH). Input resistance was determined from voltage responses evoked by hyperpolarizing current injections from -10 to -40 pA in 10-pA increments. Action potential properties were determined during depolarizing current injections of increasing amplitude (20-pA increments, 500 ms). Rheobase, $APD_{50}$, maximal depolarization and repolarization rates, amplitude, overshoot, threshold, and afterhyperpolarization were measured using the Clampfit Advanced Analysis Module.

**RNA extraction and reverse transcription from the intracardiac ganglionated region**

The intracardiac ganglionated region was microdissected, collected, and homogenized in RA1 lysis buffer containing 0.01% β-mercaptoethanol using a Polytron PT1600E tissue homogenizer. Total RNA was extracted using the NucleoSpin RNA Kit (Macherey-Nagel) according to the manufacturer's instructions. RNA concentration and purity were assessed by optical density at 260 and 280 nm using a NanoDrop ND-100 spectrophotometer.

cDNA was synthesized using random hexamer primers Pd(N)6 (Invitrogen). Total RNA was mixed with reaction buffer containing Tris-HCl, KCl, MgCl2, DTT, dNTPs, and random hexamers. RNA was denatured at 65°C for 2 min, supplemented with RNaseOUT and M-MLV reverse transcriptase, and reverse-transcribed at 37°C for 60 min. Remaining enzymes were heat-inactivated at 96°C for 2 min.

**RNA extraction and reverse transcription from isolated neurons**

After collection into patch pipettes, cell suspensions containing approximately 10 intracardiac neurons were transferred into nuclease-free microcentrifuge tubes. After centrifugation at 300 × g for 5 min, cells were resuspended in 5 μL ice-cold PBS. Cell lysis, RNA extraction, and reverse transcription were performed using the SuperScript IV CellsDirect cDNA Synthesis Kit (Invitrogen) according to the manufacturer's instructions. Negative controls included a no-cell control and a no-reverse-transcriptase control.

**Digital PCR**

Scn5a mRNA expression was quantified using a QuantStudio Absolute Q Digital PCR System (Thermo Fisher Scientific/Applied Biosystems). TaqMan Gene Expression Assays were used for Scn5a (Mm01342518_m1, FAM), ChAT (Mm01221880_m1, FAM), Tnnt2 (Mm01290256_m1, FAM), and Gapdh (Mm99999915_g1, VIC). Reaction mixtures were prepared using Absolute Q Universal DNA dPCR Master Mix (1× final concentration), TaqMan assays, and cDNA in a final volume of 10 μL.

Reaction mixtures were loaded into QuantStudio Absolute Q MAP16 plates. Each well was partitioned into 20,480 microchambers. PCR cycling consisted of enzyme activation at 96°C for 10 min, followed by 40 cycles of denaturation at 96°C for 5 s and annealing/extension at 60°C for

5 s. FAM and VIC fluorescence were acquired on the instrument and analyzed using QuantStudio Absolute Q Digital PCR Software v6. Absolute quantification was based on Poisson statistics. Results were expressed as copies/μL and normalized to Gapdh. A no-template control was included in each run.

**Supplementary Table S1. Primary and secondary antibodies**

| Antibody | Host species | Dilution | Catalog number | Supplier |
|---|---|---|---|---|
| Tyrosine hydroxylase (TH) | Rabbit | 1:300 | AB152 | Sigma-Aldrich |
| Choline acetyltransferase (ChAT) | Goat | 1:150 | AB144P | Sigma-Aldrich |
| $Na_V1.5$ | Rabbit | 1:200 | ASC005 | Alomone Labs |
| βIII-tubulin (TUJ1) | Mouse | 1:400 | T8660 | Sigma-Aldrich |
| Donkey anti-rabbit Alexa Fluor 555 | Donkey | 1:300 or 1:500 | A-31572 | Thermo Fisher Scientific |
| Donkey anti-goat Alexa Fluor 488 | Donkey | 1:300 | A-11055 | Thermo Fisher Scientific |
| Chicken anti-mouse Alexa Fluor 488 | Chicken | 1:500 | A-21200 | Thermo Fisher Scientific |

### Supplementary data

**Supplementary table S2 : Comparison of the electrophysiological properties of intracardiac neurons with an RMP < -60 mV and > -60 mV in WT and Scn5a$^{+/-}$ neurons**

| **Global** | ***Scn5a*$^{+/-}$** | **WT** |
|---|---|---|
| Capacity (pF) | 39.4 ± 5.1 (n=13) | 34.1 ± 3.4 (n=18) |
| RMP (mV) | -63.4 ± 2.4 (n=14) | -61.9 ± 2.4 (n=18) |
| Rheobase (pA) | 68.6 ± 7.2 (n=14) | 66.7 ± 9.4 (n=18) |
| Number of AP max | 2.1 ± 0.4 (n=14) | 3.6 ± 0.10 (n=18) |
| Peak amplitude (mV) | 73.6 ± 3.2 (n=14) | 79.2 ± 3.4 (n=18) |
| Threshold (mV) | -23.5 ± 1.2 (n=14) | -25.3 ± 1.1 (n=18) |
| Overshoot (mV) | 50.1 ± 2.4 (n=14) | 53.8 ± 2.5 (n=18) |
| APD at 50% | 1.8 ± 0.1 (n=14) | 1.6 ± 0.1 (n=18) |
| Rise slope | 79.5 ± 6.9 (n=14) | 99.6 ± 10.05 (n=18) |
| dV/dt max (mV/ms) | 105.9 ± 8.2 (n=14) | 128.2 ± 12.5 (n=18) |

| **RMP < -60 mV** | ***Scn5a*$^{+/-}$** | **WT** |
|---|---|---|
| Capacity (pF) | 36.4 ± 5.3 (n=8) | 38.3 ± 6.1 (n=9) |
| RMP (mV) | -70.1 ± 1.8 (n=8) | -70.9 ± 1.9 (n=9) |
| Rheobase (pA) | 65 ± 11.8 (n=8) | 62.5 ± 7.10 (n=8) |
| Number of AP max | 2.9 ± 0.6 (n=8) | 5.4 ± 1.7 (n=9) |
| Peak amplitude (mV) | **71.3 ± 4.3 (n=8)** ** | 88.8 ± 4.5 (n=9) |
| Threshold (mV) | **-24.0 ± 1.8 (n=8)** * | -28.6 ± 1.1 (n=9) |
| Overshoot (mV) | **47.3 ± 29 (n=8)** ** | 60.2 ± 3.7 (n=9) |
| APD at 50% | **1.7 ± 0.1 (n=8)** * | 1.4 ± 0.1 (n=9) |
| Rise slope | **80.8 ± 10.8 (n=8)** * | 127.6 ± 12.6 (n=9) |
| dV/dt max (mV/ms) | **109.1 ± 13.2 (n=8)** * | 162.8 ± 16.1 (n=9) |

| **RMP > -60 mV** | ***Scn5a*$^{+/-}$** | **WT** |
|---|---|---|
| Capacity (pF) | 44.2 ± 10.6 (n=5) | 29.9 ± 2.6 (n=9) |
| RMP (mV) | -54.5 ± 1.3 (n=6) | -52.8 ± 1.1 (n=9) |
| Rheobase (pA) | **73.3 ± 6.7 (n=6)** * | 55.5 ± 8.0 (n=9) |
| Number of AP max | 1 ± 0 (n=6) | 1.8 ± 0.4 (n=9) |
| Peak amplitude (mV) | 76.7 ± 5.1 (n=6) | 69.5 ± 2.4 (n=9) |
| Threshold (mV) | -22.8 ± 1.4 (n=6) | -22.1 ± 1.3 (n=9) |
| Overshoot (mV) | 53.9 ± 3.9 (n=6) | 47.5 ± 1.9 (n=9) |
| APD at 50% | 1.9 ± 0.1 (n=6) | 1.8 ± 0.1 (n=9) |
| Rise slope | 77.8 ± 8.2 (n=6) | 71.6 ± 8.6 (n=9) |
| dV/dt max (mV/ms) | 101.7 ± 9.1(n=6) | 93.6 ± 10.5 (n=9) |

dV/dt : maximum rate of the depolarization or repolarization phase. AHP duration and amplitude were only calculated for neuron displaying AHP. RMP : Resting membrane potential (mV). *P<0.05 ; **P<0.01 Mann Whitney test.
